# Biomolecular recognition of the glycan neoantigen CA19-9 by distinct antibodies

**DOI:** 10.1101/2021.02.17.431565

**Authors:** Aliza Borenstein-Katz, Shira Warszawski, Ron Amon, Nova Tasnima, Hai Yu, Xi Chen, Vered Padler-Karavani, Sarel Jacob Fleishman, Ron Diskin

**Affiliations:** Department of Chemical and Structural Biology, Weizmann Institute of Science, 76100 Rehovot, Israel; Department of Biomolecular Sciences, Weizmann Institute of Science, 76100 Rehovot, Israel; Department of Cell Research and Immunology, The Shmunis School of Biomedicine and Cancer Research, The George S. Wise Faculty of Life Sciences, Tel Aviv University, Tel Aviv 69978, Israel; Department of Chemistry, University of California, Davis, CA 95616, USA

## Abstract

Glycans decorate cell surface, secreted glycoproteins and glycolipids. Altered glycans are often found in cancers. Despite their high diagnostic and therapeutic potentials, glycans are polar and flexible molecules that are quite challenging for the development and design of high-affinity binding antibodies. To understand the mechanisms by which glycan neoantigens are specifically recognized by antibodies, we analyze the biomolecular recognition of a single tumor-associated carbohydrate antigen CA19-9 by two distinct antibodies using X-ray crystallography. Despite the plasticity of glycans and the very different antigen-binding surfaces presented by the antibodies, both structures reveal an essentially identical extended CA19-9 conformer, suggesting that this conformer’s stability selects the antibodies. Starting from the bound structure of one of the antibodies, we use the AbLIFT computational method to design a variant with seven core mutations that exhibited tenfold improved affinity for CA19-9. The results reveal strategies used by antibodies to specifically recognize glycan antigens and show how automated antibody-optimization methods may be used to enhance the clinical potential of existing antibodies.

## INTRODUCTION

Cancer is a leading cause of death worldwide, with epithelial carcinoma the most devastating. Changes in cell surface markers are one of the hallmarks of cancer, and antibodies that bind these markers are ideal therapeutics and/or diagnostic tools [1]. Surface glycosylation is a universal feature of cells but is often altered during malignant transformation, leading to a distinct subset of antigens that are selectively and abundantly expressed on cancer cells [2–5]. This feature is intimately associated with abnormal expression of the glycosylation biosynthetic pathways, leading to variations in the basic core carbohydrate chains (glycans) conjugated to glycoproteins and glycolipids [3,6]. These aberrations particularly affect the expression of sialic acids (Sias) that cap cell surface glycans. For example, the sialyl Lewis a (SLe^a^) tetrasaccharide stems from incomplete synthesis of the normal glycan disialyl LeA. While both SLe^a^ and disialyl Le^a^ are generated via the same metabolic pathway, reduction or loss of expression of the α2–6-sialyltransferase (ST6GalNAc VI) during malignancy shifts the pathway towards expression of the cancer antigen SLe^a^ [7] (Fig. 1). Altered glycosylation pattern often correlate with advanced cancer stage, progression and/or metastasis [2,4,5,8].

**Fig. 1.**
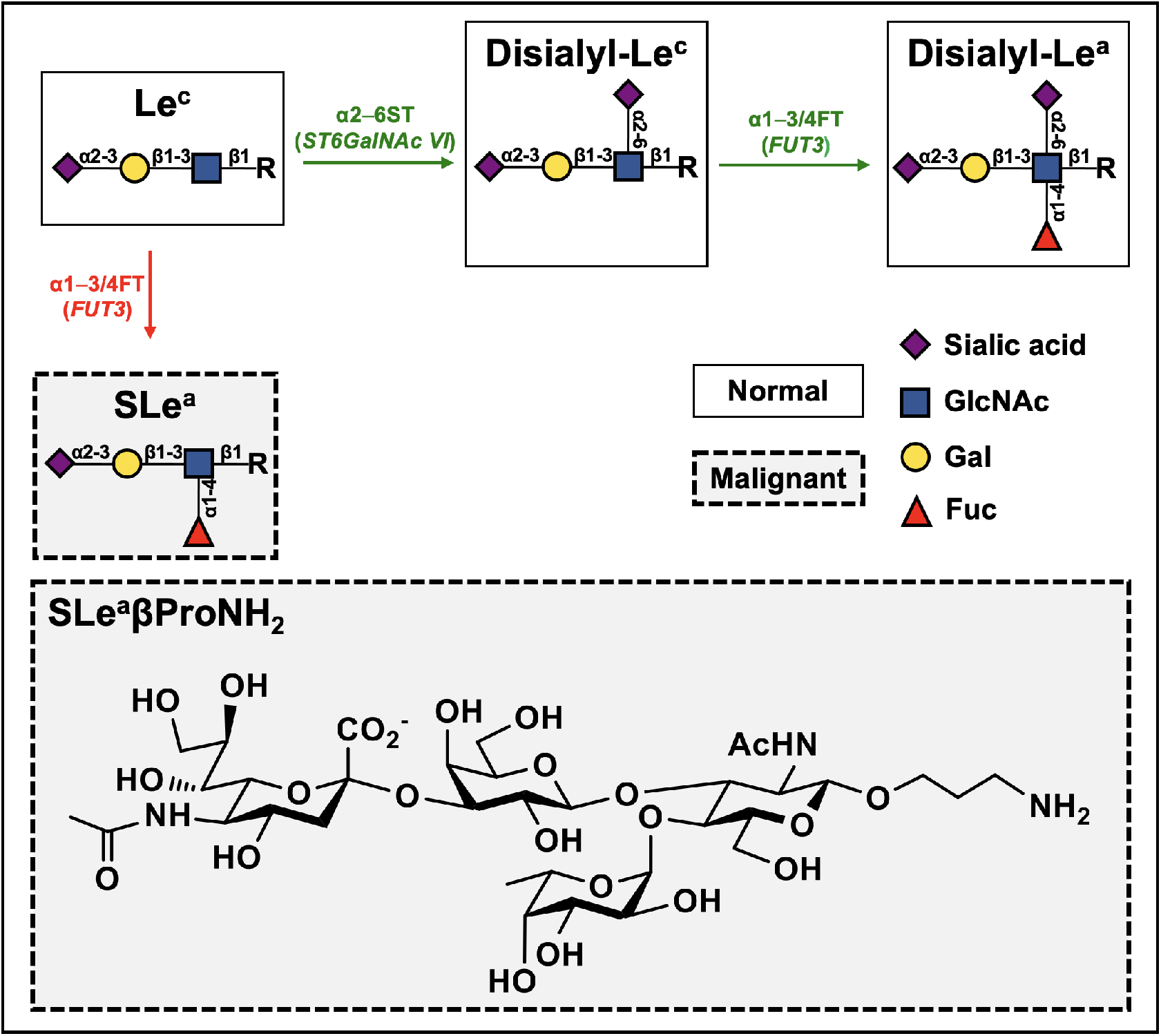
Biosynthetic pathway of SLe^a^ and Disialyl-Le^a^. SLe^a^ (CA19-9) is a Type-1 tetrasaccharide tumor-associated carbohydrate antigen composed of fucose (Fuc), *N-*acetylglucosamine (GlcNAc), galactose (Gal), and sialic acid (Sia). In the normal biosynthetic pathway, the precursor Le^c^ is commonly further elongated by α2–6-sialyltransferase and α1–3/4-fucosyltransferase to generate disialyl-Le^a^, which has an additional sialic acid moiety compared to SLe^a^. The SLe^a^βProNH_2_ probe, Neu5Acα2–3Galβ1–3(Fucα1–4)GlcNAcβO(CH_2_)2CH_2_NH_2_, is a SLe^a^ antigen with a linker containing a terminal primary amine which can be used for conjugation for functional studies.

SLe^a^, also known as carbohydrate antigen CA19-9, is detected on pancreatic, colorectal, stomach and liver cancers [7,9]. This cancer-associated marker is widely used in clinical practice for serological assays [5,10,11]. It is the only FDA-approved test for pancreatic cancer and is also used in assays for colorectal, gastric and biliary cancers [5]. The assay is based on a monoclonal antibody (mAb) capturing the CA19-9 antigen, and is commonly used to monitor clinical response to therapy; however, it is not useful for early detection or diagnosis due to both false positive and false negative readouts [10–12]. While this serological assay has been available for almost three decades, the interpretation of CA19-9 measurements is largely hampered by nonspecific increase reads for the levels of CA19-9, either due to associated morbidity (e.g. obstruction of the biliary tree or inflammation) or due to assay-dependent variability, both in diseased and healthy subjects [13]. As a result, pancreatic cancer is often detected too late at an advanced stage resulting in a very short five-year survival rate. Interestingly, a recent study in mice demonstrated that CA19-9 is an active driver of pancreatitis, which leads to the development of pancreatic cancer [14]. This discovery assigns, for the first time, an active role for CA19-9 as a cancer driver. Importantly, mAbs targeting CA19-9 were able to reverse pancreatitis in this mouse model [14], establishing CA19-9 as a prime target for cancer therapy.

A potential obstacle for using anti-carbohydrate antibodies for theranostics is their low affinity and low specificity compared to antibodies targeting proteins [15,16]. This limitation prompted development of tools to better define such antibody-antigen interactions [17] and enhance their affinity [18]. Thus, detailed structural information for the CA19-9 and its recognition by mAbs is a step towards the design of more efficient reagents in the fight against some of the most devastating cancer types. Recently, we described a new antibody-design method, called AbLIFT, that focuses design calculations on the interfaces formed between specific antibody light and heavy chain pairs as observed in crystallographic analysis [19]. AbLIFT optimizes the computed energy of a variety of antibodies. Furthermore, in experimental screens of up to 20 designed antibodies, several of the designs had increased expressibility, thermal stability, aggregation resistance and antigen-binding affinity [19]. Thus, we hypothesized that AbLIFT could also be used to enhance anti-CA19-9 antibodies.

Here, we provide molecular insights into antigen recognition by two of the most well-defined anti-CA19-9 mAbs, the murine 1116NS19.9 [20,21] commonly used in the CA19-9 serological test, and the human 5b1 [22] that is currently investigated for cancer imaging in clinical trials [23]. We present high-resolution crystal structures of both antibodies in complex with CA19-9 antigen. These structures reveal two distinct binding solutions to a single conformer of CA19-9. We further use the state-of-the-art AbLIFT computational tool together with this structural information to design mAbs that target CA19-9 with an order of magnitude greater affinity.

## RESULTS

### Cloned 1116NS19.9 and 5b1 are specific to SLe^a^

To reveal the molecular basis for the immune recognition of the tumor-associated carbohydrate antigen CA19-9, we selected for structural studies two of the most widely used mAbs, 1116NS19.9 and 5b1. The variable domains of the mAbs were cloned and their functionality were confirmed both as single-chain Fv (scFv) fragments, and as full-length human IgG1. First, sequences of the variable heavy chain (V_H_) and of the variable light chain (V_L_) fragments of 1116-NS-19-9 and 5b1 were each cloned into pETCON2 plasmid as scFv with (G4S)3 linker between the V_H_ and the V_L_ domains, and were transformed to yeast cells which were induced to express cell surface scFvs [24]. Antigen recognition was evaluated in solution by flow cytometry of scFv-expressing yeast cells against SLe^a^-nanoparticles with multivalent expression of antigen to resemble their presentation on cancer cells. This analysis revealed similar scFv surface expression, however better antigen binding with 5b1-scFv than with 1116NS19.9-scFv (Fig. 2a). Subsequently, sequences encoding the variable regions of the two mAbs were cloned into a human IgG1 scaffold and antigen recognition was examined. Binding of full-length antibodies to multivalent-nanoparticles coated on a solid surface showed binding to SLe^a^, but complete loss of binding to Le^a^ antigen that lacks the terminal sialic acid (Fig. 2b), implying that sialic acid recognition plays an important role for the binding of both antibodies. These data also show that the cloned antibodies are fully functional, both against a flexible antigen in solution and with antigen fixed to a solid surface, and are specific for their target antigen SLe^a^ (CA19-9).

**Fig. 2.**
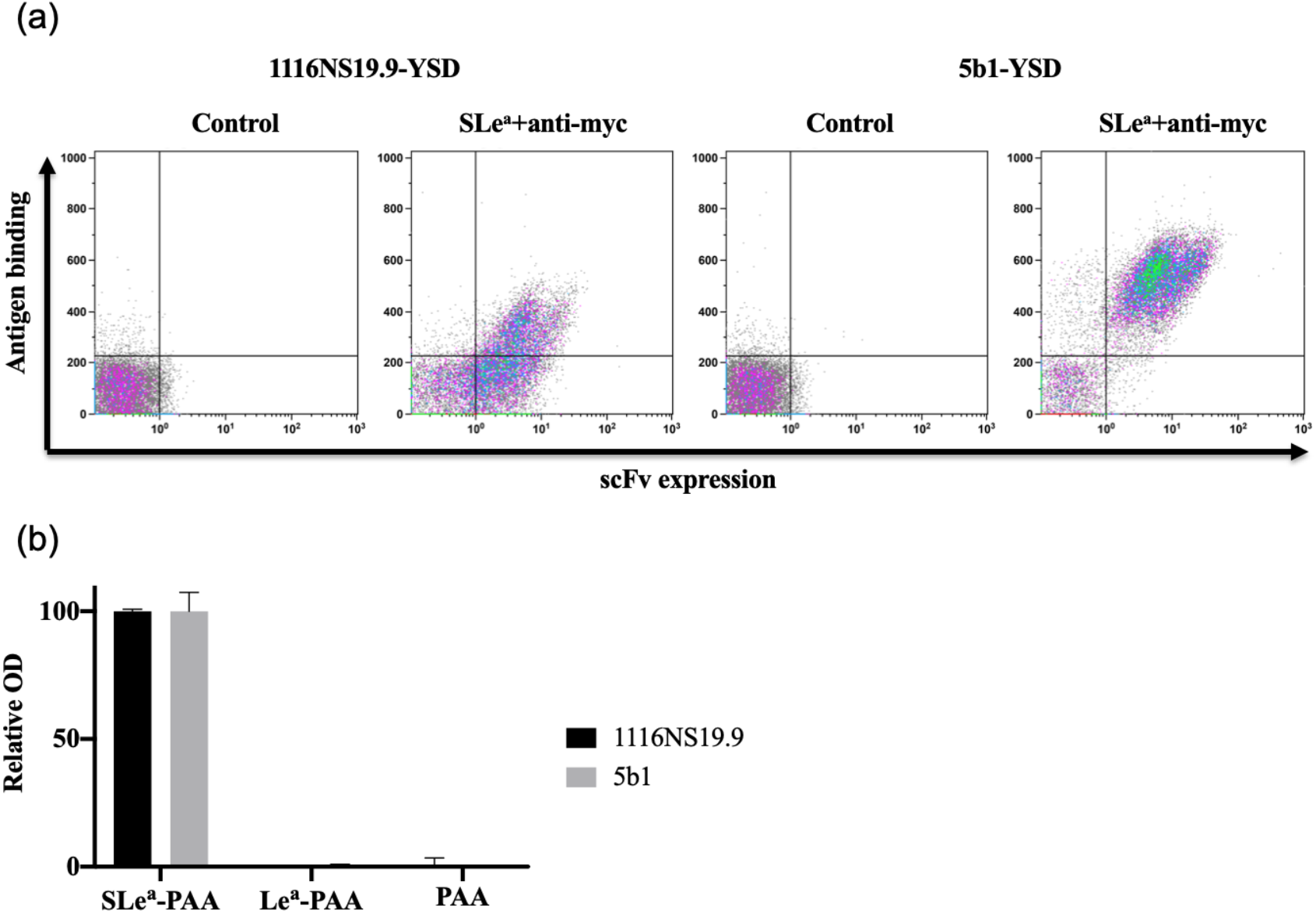
Expression of 1116NS19.9 and 5b1 antibodies and their antigen recognition. (a) FACS analyses of yeast cells expressing either 1116NS19.9, 5b1, or an empty vector as negative controls. The X-axis indicates the surface display levels (by detecting C’-c-Myc tag) and the Y-axis indicates antigen binding (using 1 μM SLe^a^-PAA-Bio – multivalent SLe^a^-polyacrylamide polymers tagged with biotin). These are representative plots from at least two independent repeats. (b) Specificity of antigen binding. Binding of 1116NS19.9 and 5b1 in IgG forms to solid surface coated with SLe^a^-PAA-Bio, Le^a^-PAA-Bio, and PAA-Bio was evaluated by ELISA in duplicates. Relative optical density (OD) was calculated as percentage of maximal binding of each antibody clone (mean ± SEM; Representative of at least two independent repeats).

### Determining the CA19-9-bound structures of the two mAbs

As a first step towards determining the molecular structure of the two antibodies bound to CA19-9, we produced the two mAbs in HEK293F cells and purified them using protein-A affinity chromatography. Fab fragments were obtained using a papain digest of the IgGs followed by separation of the Fabs from the Fc fragments using protein-A affinity and size exclusion chromatography (Fig. 3a). CA19-9 (SLe^a^) antigen was chemoenzymatically synthesized as described in ref. [25], with a terminal primary amine-containing linker (SLe^a^βProNH_2_; Fig. 1). We crystallized both the apo (without antigen) and the holo (with CA19-9 antigen) states of Fab fragments of 1116NS19.9 and 5b1 mAbs, followed by X-ray diffraction analyses at the European Synchrotron Radiation Facility. Complete data sets were collected at 1.6 Å and 1.5 Å resolutions for the CA19-9-bound and the apo-state of mAb 1116NS19.9, respectively (Table 1). Complete data sets were further collected at 2.4 Å resolutions for the CA19-9-bound and the apo-Fab of Ab 5b1 (Table 1), and all structures were solved using molecular replacement. For the apo mAb 1116NS19.9, a human-derived Fab (PDB: 3U7W) was used as the search model and subsequently the 1116NS19.9 structure was used as a search model for solving the rest of the structure through molecular replacement. In the holo-structures of both 1116NS19.9 and 5b1, clear electron density for CA19-9 was observed, allowing to accurately model it (Fig. 3a). In both structures, density for the propyl linker attached to the CA19-9 was either missing completely (5b1) or was only weakly visible for the first carbon atom of the linker (1116NS19.9). Hence this linker was omitted from the models.

**Fig. 3.**
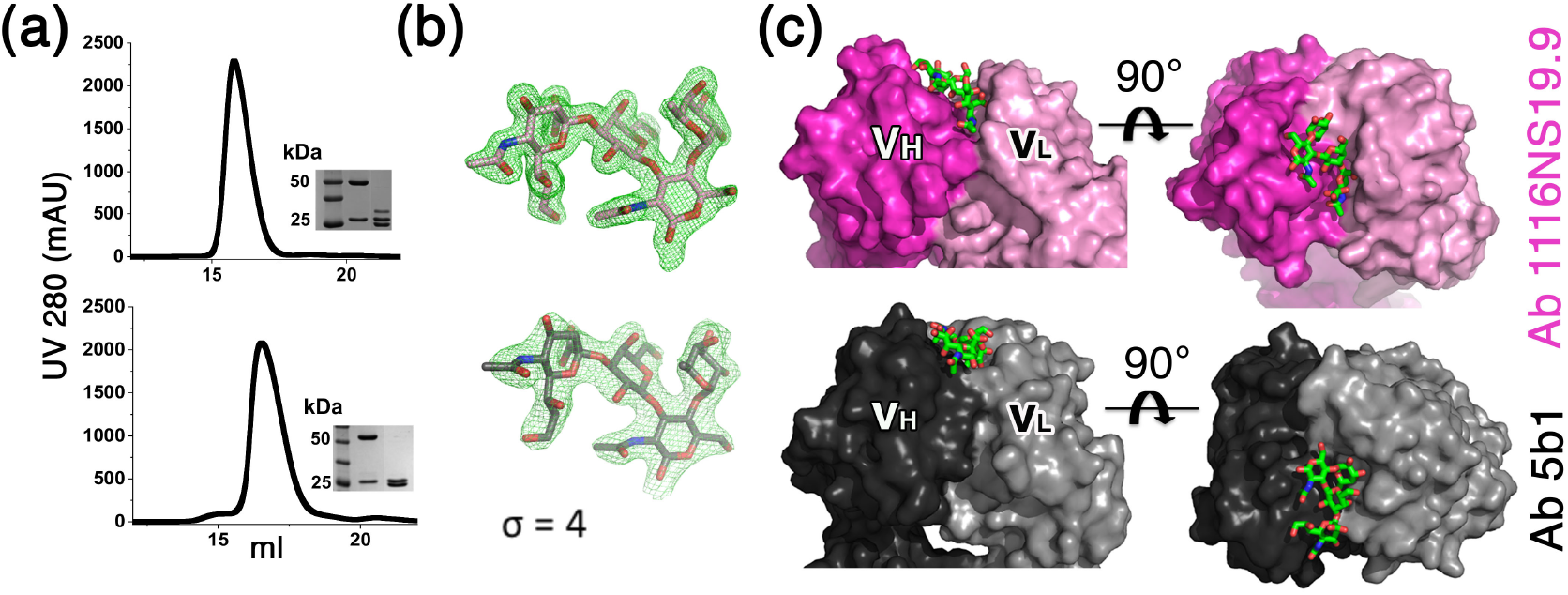
Holo-structure of 1116NS19.9 and 5b1 Fabs. (a) Production and purification of Fab fragments. Chromatograms for the Fab fragments of 1116NS19.9 (top) and 5b1 (bottom) indicate their migration profile in the size-exclusion chromatography column. Images of Coomassie-stained SDS-PAGE are shown in the insets, loaded with a size marker, the IgG fraction, and the cleaved Fab fraction (left to right). (b) CA19-9 in Fo-Fc (difference) omit maps. CA19-9 structures from 1116NS19.9 (top) and from 5b1 (bottom) are shown in Fo-Fc maps (green mesh) at σ=4. These difference maps were calculated from the final models after omitting the CA19-9 molecules. (c) The binding pockets for CA19-9. The variable domains of 1116NS19.9 and of 5b1 are shown using surface representations. The chains are colored in dark pink and light pink for the heavy and light chains of 1116NS19.9, respectively, and in dark gray and light gray for the heavy and light chain of 5b1. The CA19-9 molecules are shown using stick representation.

**Table 1:** Data collection and refinement statistics.

| Parameter | Holo<br>1116NS19.9 | Apo<br>1116NS19.9 | Holo 5b1 | Apo 5b1 | AbLIFT-15 |
| --- | --- | --- | --- | --- | --- |
| PDB ID | 6XTG | 6XUD | 6XUN | 6XUL | 6XUK |
| Wavelength (Å) | 0.97242 | 0.97242 | 0.87313 | 0.87313 | 0.97371 |
| Space group | $P 2_1 2_1 2$ | $P 4_1 2_1 2$ | $P 3_1$ | $P 3_2$ | $P 4_3 2_1 2$ |
| Cell dimensions |  |  |  |  |  |
| a, b, c (Å) | 89.17 60.20<br>84.44 | 64.92 64.92<br>242.6 | 152.54 152.54<br>60.89 | 155.01 155.01<br>121.78 | 64.93 64.93<br>244.9 |
| $\alpha, \beta, \gamma$ ° | 90 90 90 | 90 90 90 | 90 90 120 | 90 90 120 | 90 90 90 |
| Resolution (Å) | 44.58-1.55<br>(1.60-1.55)<sup>a</sup> | 45.9-1.51<br>(1.56-1.51)<sup>a</sup> | 47.58-2.41<br>(2.48-2.41)<sup>a</sup> | 46.84-2.41<br>(2.50-2.41)<sup>a</sup> | 62.75-1.42<br>(1.47-1.42)<sup>a</sup> |
| $R_{\text{meas}}$ (%) | 5.9 (158)<sup>a</sup> | 13.4 (103.1)<sup>a</sup> | 17.42 (90.17)<sup>a</sup> | 19.24 (103.8)<sup>a</sup> | 10.85 (111.8)<sup>a</sup> |
| $CC_{1/2}$ | 99.9 (56.8)<sup>a</sup> | 99.6 (79.2)<sup>a</sup> | 99.1 (67.9)<sup>a</sup> | 98.3 (53.3)<sup>a</sup> | 99.6 (57.7)<sup>a</sup> |
| $I/\sigma I$ | 13 (1.1)<sup>a</sup> | 9.4 (2.3)<sup>a</sup> | 6.62 (1.81)<sup>a</sup> | 5.58 (1.35)<sup>a</sup> | 9.4 (2.3)<sup>a</sup> |
| Completeness (%) | 95 (95.2)<sup>a</sup> | 99.9 (100)<sup>a</sup> | 99.76 (98.2)<sup>a</sup> | 99.70 (99.9)<sup>a</sup> | 99.89 (99.6)<sup>a</sup> |
| Multiplicity | 2.9 (3)<sup>a</sup> | 9.1 (9.4)<sup>a</sup> | 5.1 (5.2)<sup>a</sup> | 3.5 (3.5)<sup>a</sup> | 6.1 (5.6)<sup>a</sup> |
| Reflections | 359539 | 753716 | 309763 | 446316 | 610565 |
| Unique reflections | 64876 | 82652 | 61168 | 126376 | 99646 |
| Refinement |  |  |  |  |  |
| Resolution (Å) | 44.58 – 1.55 | 45.32 – 1.51 | 47.58 – 2.41 | 46.84 – 2.41 | 62.75 – 1.42 |
| No. of reflections | 64837 | 82525 | 61030 | 126037 | 99619 |
| $R_{\text{work}}/R_{\text{free}}$ (%) | 17.39 / 20.13 | 14.69 / 17.32 | 16.99 / 21.34 | 16.78 / 22.95 | 14.9 / 18.49 |
| No. of atoms |  |  |  |  |  |
| Protein | 3356 | 3361 | 9978 | 19904 | 3312 |
| Ligand/ion | 57 | - | 192 | - | 68 |
| Water | 382 | 643 | 485 | 1022 | 661 |
| B factors |  |  |  |  |  |
| Protein | 38.67 | 21.43 | 45.35 | 49.5 | 20.16 |
| Ligand/ion | 33.85 | - | 44.92 | - | 23.93 |
| Water | 45.09 | 35.81 | 44.41 | 46.33 | 34.20 |
| Ramachandran |  |  |  |  |  |
| Favored (%) | 98.12 | 98.83 | 97.60 | 96.54 | 98.12 |
| Allowed (%) | 1.88 | 1.17 | 2.32 | 3.34 | 1.41 |
| Outlier (%) | 0.0 | 0.0 | 0.08 | 0.12 | 0.47 |
| RMSD |  |  |  |  |  |
| Bond length (Å) | 0.006 | 0.008 | 0.006 | 0.008 | 0.013 |
| Bond angles ° | 0.733 | 0.995 | 1.17 | 1.22 | 1.65 |

<sup>a</sup> Values in parentheses are for the highest resolution-shell

### The two antibodies recognize a similar, low-energy conformer of CA19-9

The structures of mAbs 1116NS19.9 and 5b1 reveal that in both cases, the CA19-9 antigen binds in a groove that is formed between the variable heavy (V_H_) and variable light (V_L_) domains (Fig. 3c). Typically, glycosidic bonds can freely rotate, and hence oligosaccharides inherently present an ensemble of conformations in solution [26]. Nevertheless, comparing the bound CA19-9 antigen from the structures of 1116NS19.9 and of 5b1 reveals that in both cases, CA19-9 assumes a surprisingly similar conformer (Fig. 4a) with an all-atom RMSD (root-mean-square deviation) of only 0.46 Å. Of note, 1116NS19.9 and 5b1 were isolated from different species (i.e., human vs. mouse) and were elicited against CA19-9 that was displayed in two very distinct contexts (i.e., on a carcinoma cell and as a protein-conjugated antigen). The fact that both antibodies have evolved to recognize a similar conformation of CA19-9 implies that the conformer observed in our crystallographic analyses is energetically preferred.

**Fig 4.**
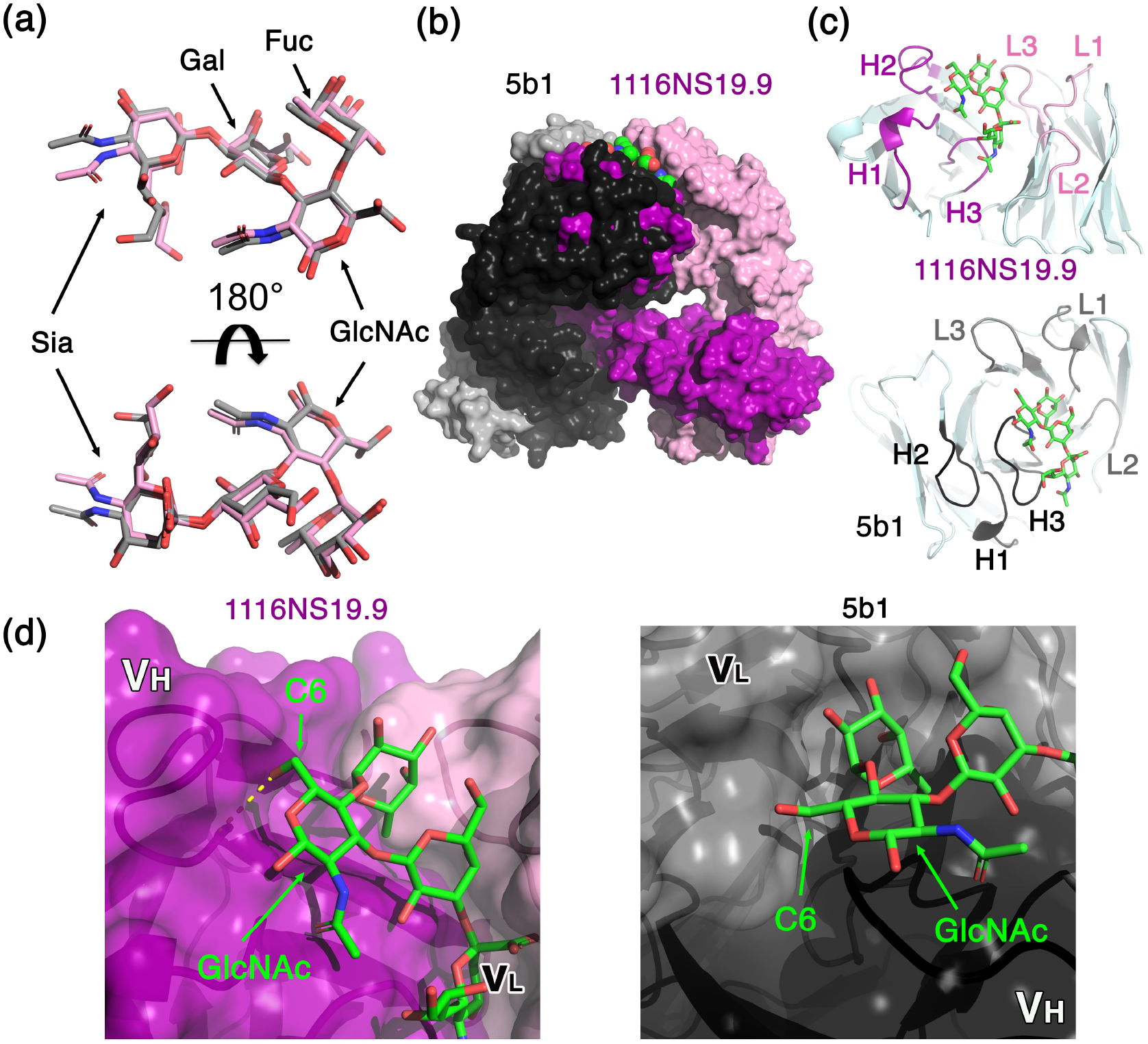
Molecular recognition of a similar, extended, CA19-9 conformer. (a) CA19-9 assumes a similar extended conformation in both structures. Superimposition of CA19-9 from the 1116NS19.9 structure (pink) and of CA19-9 from the 5b1 structure (gray) is shown in two different orientations, as indicated. The saccharides labeled are L-fucose (Fuc), *N*-acetylglucosamine (GlcNAc), galactose (Gal), and sialic acid (Sia) (b) The relative binding orientations of 1116NS19.9 and 5b1 to CA19-9. The two structures are superimposed based on the CA19-9 antigen. 5b1 and 1116NS19.9 are shown in gray and pink surfaces, using dark and light colors for the heavy and light chains, respectively. (c) The CDRs’ role in binding CA19-9. Ribbon diagrams of 1116NS19.9 (top) and 5b1 (bottom) illustrate the organization of the CDRs around CA19-9, which is shown in the same orientation in both images. CDRs are highlighted and labeled. (d) Molecular basis for CA19-9 selectivity. Surface representations of 1116NS19.9 (left) and 5b1 (right) show the C6 position of the GlcNAc. The hydrogen bond that the C6 hydroxyl is forming with 1116NS19.9 is indicated with a dashed yellow line.

The unanticipated finding that the two mAbs recognize a similar conformer of CA19-9 raises the question of whether they also utilize similar binding mechanisms. By superimposing the structures according to the CA19-9 antigen, it is clear that the antibodies bind CA19-9 in two distinct ways (Fig. 4b). The two Fabs approach CA19-9 from different angles (Fig. 4b) and use their complementarity-determining regions (CDRs) for binding in different modes (Fig. 4c). In the case of mAb 1116NS19.9, aside from CDRL1, all other CDRs are involved in ligand binding, and CDRL3 is the most significant contributor to molecular interactions. By contrast, mAb 5b1 does not engage CA19-9 through CDRH1 and CDRH2, while CDRH3 and CDRL1 make significant contacts with CA19-9 (Fig. 4c). Comparing the binding of CA19-9 by 1116NS19.9 to 5b1, the former has a relatively deeper groove than the latter, which is reflected in a larger total buried surface area for the complex formation (973 and 859 Å^2^ for 1116NS19.9 and 5b1, respectively), though both antibodies exhibit a similar binding specificity profile for CA19-9.

### Molecular basis for the specificity toward CA19-9

CA19-9 differs from disialyl-Le^a^ antigen by the lack of a sialic-acid (Sia) moiety that is typically connected to the carbon-6 (C6) of the GlcNAc (Fig. 1). In the case of 1116NS19.9, the hydroxyl group extending from C6 directly faces the heavy-chain, leaving no room for accommodating the extra Sia in this conformation (Fig. 3d). Moreover, the binding of CA19-9 to 1116NS19.9 is partly facilitated by a hydrogen bond that the GlcNAc C6-hydroxyl group forms with Asn52A on CDRL2, favoring a free hydroxyl group in this position. In the case of mAb 5b1, the hydroxyl extension from C6 of the GlcNAc is not buried at the interface, as seen with 1116NS19.9 (Fig. 4d). Nevertheless, an additional Sia moiety cannot be accommodated unless the Sia assumes a conformation that is potentially less energetically favorable than the one observed in both antibody-bound complexes. Moreover, the free hydroxyl of C6 participates in a water-mediated interaction with the light-chain of 5b1, providing an additional selection power for CA19-9 in this conformation.

### Molecular recognition of CA19-9

Considering the relatively small size of CA19-9 (819 Dalton) and its hydrophilic nature, it could be regarded as a suboptimal immunogen. As such, the recognition of CA19-9 by the mAbs could be suboptimal and restricted to only a few contacts. Nevertheless, 1116NS19.9 and 5b1 display intricate interaction networks with CA19-9 (Fig. 5a & 5b) that are likely to be critical for binding affinity and specificity to this challenging antigen. In the case of 1116NS19.9, all the hydroxyl groups of CA19-9 that face the antibody, aside from O7 of the sialic acid’s hydroxyl, participate in either direct or water-mediated polar interactions with the mAb (Fig. 5a). A large number of residues from both chains mediate polar contacts with the antigen, including heavy chain positions Trp33, Asn52A, Arg95, and Phe96, as well as the light-chain positions Tyr49, Arg50, Arg53, Tyr91, Asp92, and Arg96, mediate these polar interactions (Fig. 5a). Of these interactions, the guanidino group of the light-chain Arg50 is especially important as it forms an ideal salt-bridge with the carboxylate of the Sia (Fig. 5a). A few additional hydrophobic interactions contribute to and complete the recognition site of CA19-9 on 1116NS19.9 (Fig. 5a). In the case of 5b1, there is also a saturated network of polar interactions that include all the hydroxyls of CA19-9 that face the antibody, aside from O4 of the Gal (Fig. 5b); some of these interactions are water-mediated. On the heavy-chain, Arg97, Arg98, Thr100A, Gly100B, and Ala100D, all from CDRH3, contact CA19-9 (Fig. 5b). The light-chain residues Ser30B, Phe32, Tyr34, Arg50, Trp50, Trp91, and Asp93 complete the set of residues that form the polar binding site of CA19-9. Interestingly, the carboxyl group of the Sia in the case of 5b1 is exposed to the solvent and does not contribute to the binding of CA19-9. In addition, and similarly to 1116NS19.9, several hydrophobic interactions contribute to the binding of the CA19-9 antigen (Fig. 5b).

**Fig 5.**
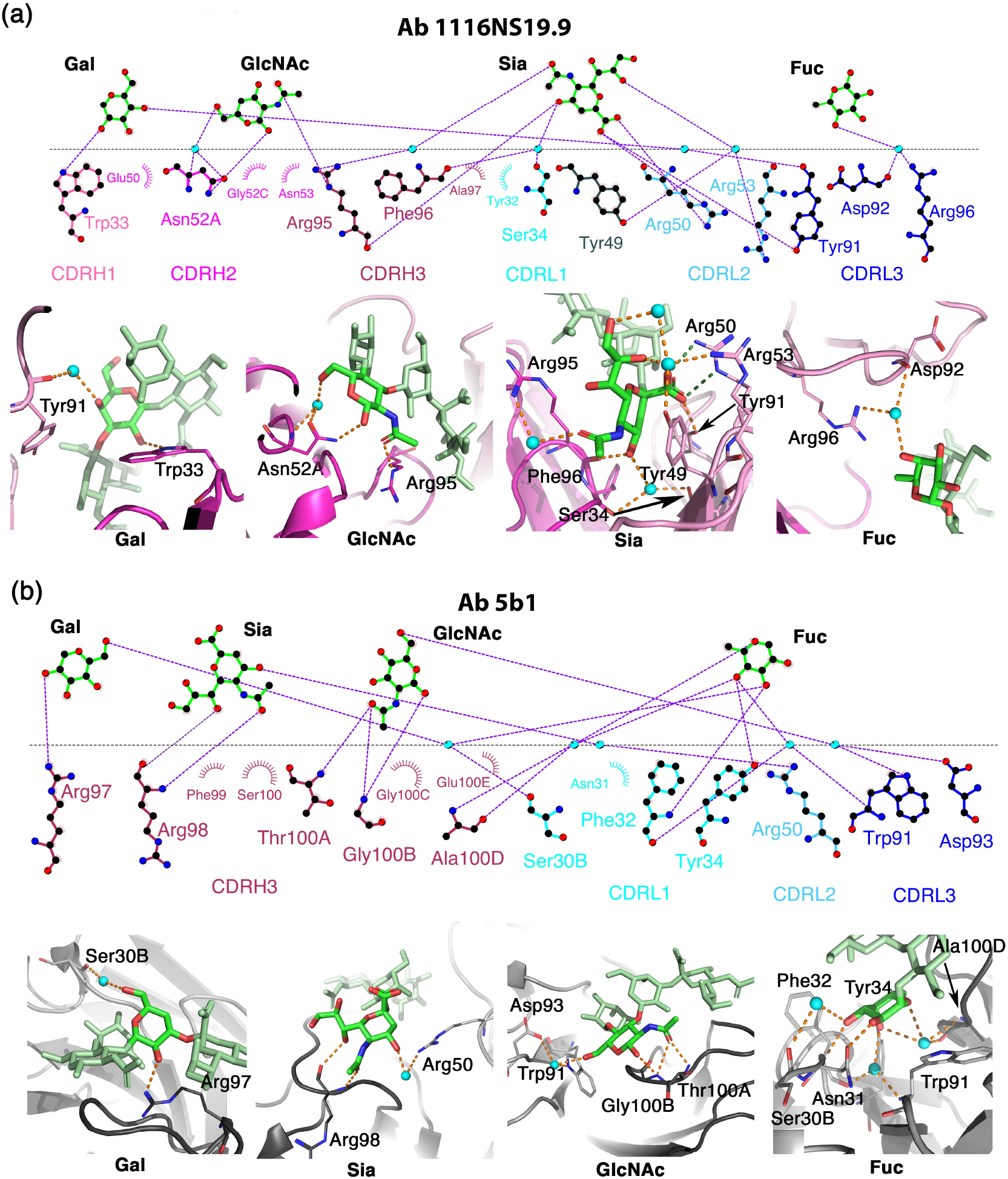
The hydrophilic binding sites of CA19-9. (a) An overview of the CA19-9 binding site on 1116NS19.9. The saccharides are labeled L-fucose (Fuc), *N*-acetylglucosamine (GlcNAc), galactose (Gal), and sialic acid (Sia) (b) An overview of the CA19-9 binding site on 5b1. For both 1116NS19.9 and 5b1, upper panels show 2D representations of the interactions between CA19-9 and the antibodies that were generated using LIGPLOT [27]. The monosaccharide moieties of CA19-9 were separated for clarity. Water molecules are indicated with spheres in cyan, and polar bonds are indicated with dashed purple lines. CDRs are labeled and differentially colored. In the lower panels, the interactions of the antibodies with each of the sugar moieties of CA19-9 are shown. The heavy and light chains are colored in purple/pink and black/grey for 1116NS19.9 and 5b1, respectively. Water molecules are indicated with spheres in cyan. Orange and green dashed lines indicate hydrogen bonds and salt-bridge interactions, respectively.

### Binding-induced conformational changes

In order to bind CA19-9, 1116NS19.9 and 5b1 may need to undergo some conformational changes. To evaluate this possibility, we compared the apo and the holo-structures of 1116NS19.9 (Fig. 6a) and of 5b1 (Fig. 6b). Indeed, the superimposition of the apo and holo 1116NS19.9 reveals some significant conformational changes between the structures (Fig. 6a). Trp33, together with Arg95 from CDRH1 and CDRH3, respectively, form a tight cation-π interaction in the apostate of 1116NS19.9. For binding CA19-9 in the holo-structure, this cation-π interaction breaks, and both residues assume different rotamers that allow them to accommodate and to engage with CA19-9 through hydrogen bonds (Fig. 6a). Also, Asn53 of CDRH2 assumes a different rotamer that allows the entire loop to move closer to CA19-9, which enables Asn52A to form a hydrogen bond with the hydroxyl of the GlcNAc-C6 (Fig. 6a & 4d). In the light chain of 1116NS19.9, Arg50 of CDRL2 forms cation-π interaction with a nearby tyrosine, and upon binding to CA19-9, this interaction breaks, and Arg50 assumes a different rotamer that allows it to form a salt-bridge with the carboxyl group of the Sia moiety (Fig. 6a). In contrast to these substantial structural rearrangements of 1116NS19.9, the superimposition of the apo and holo-structures of 5b1 reveals that the binding of CA19-9 does not involve any notable conformational changes (Fig. 6b). Hence, not only do these antibodies recognize CA19-9 at different angles (Fig. 4b), they also evolved to use different binding mechanisms. The binding site of 5b1 is pre-configured for binding, whereas the conformational changes of 1116NS19.9 that are coupled to the abolishment of strong cation-π interactions suggest an induced-fit mechanism.

**Fig 6.**
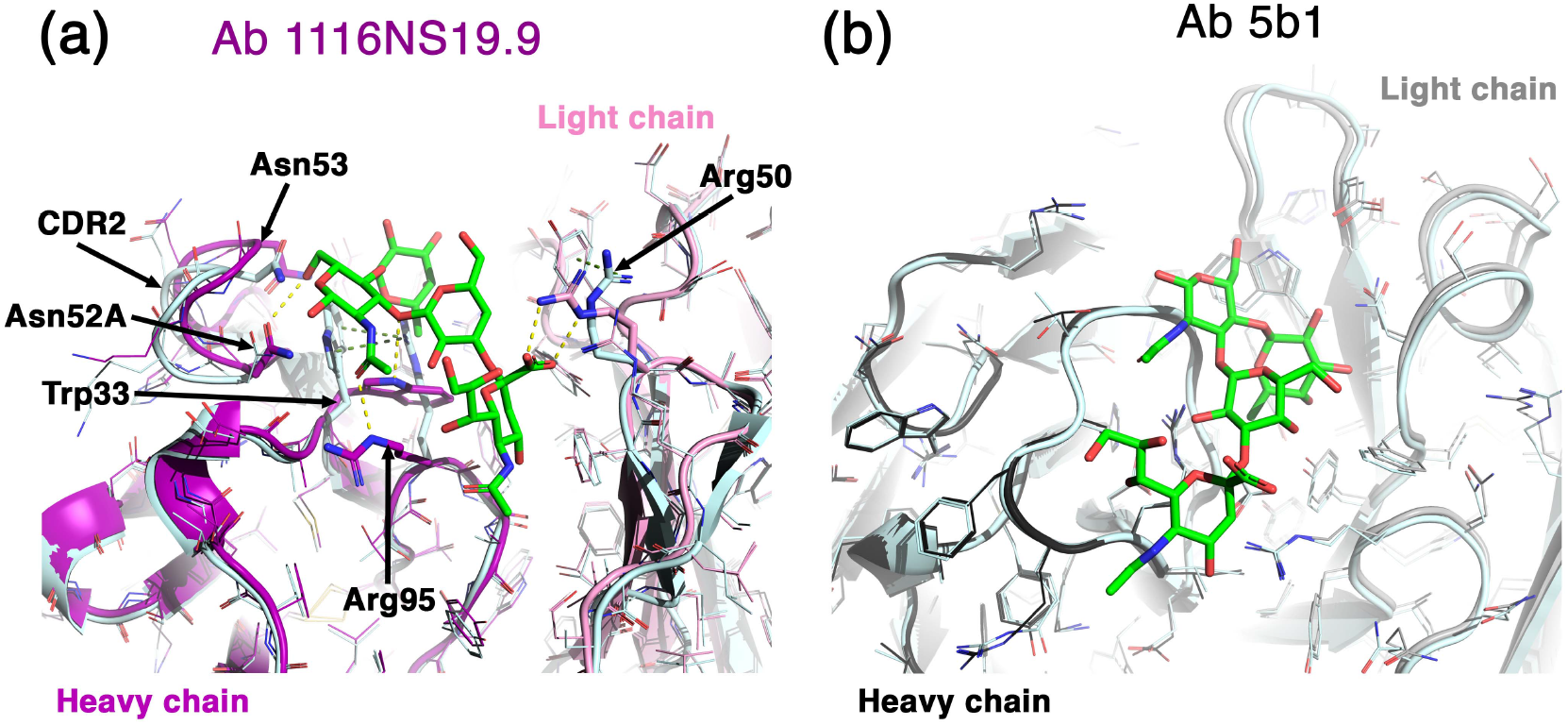
Conformational changes following binding to CA19-9. (a) Superimposition of apo 1116NS19.9 (pale cyan) and CA19-9-bound holo 1116NS19.9 (purple and pink for the heavy and light chains, respectively). Important residues that change conformation are highlighted as sticks. Hydrogen and cation-π bonds are indicated with dashed yellow and green lines, respectively. (b) Superimposition of apo 5b1 (pale cyan) and CA19-9-bound holo 5b1 (black and grey for the heavy and light chains, respectively).

### Enhancing 1116NS19.9 affinity using computational antibody design

Increasing sensitivity of CA19-9 recognition is of great interest for advancing diagnosis and management of CA19-9-positive malignancies in general, and of pancreatic cancer in particular. Focusing on the clinically used 1116NS19.9, we sought to enhance its affinity toward CA19-9. Conventional approaches to design or engineer higher-affinity antibody variants direct efforts to alter the antigen-binding surface [24]. The intricate and highly polar antigen-binding surface revealed by the crystallographic analyses of the two antibody-bound complexes suggested, however, that mutations at these surfaces might destabilize the complex. By contrast, the recently published AbLIFT computational antibody-design method focuses design calculations on the interfaces formed between the antibody light and heavy chains, away from the antigen-binding surface (available as a web server for academic users http://AbLIFT.weizmann.ac.il) [19]. While these interfaces are often tightly packed and therefore challenging for computational or experimental strategies, optimizing them provides the important advantage that it carries a lower risk of altering the critical antibody-antigen interactions; these interactions, as revealed by our crystallographic analysis, are remarkably intricate in this particular case. Furthermore, based on the induced-fit mechanism revealed for 1116NS19.9 recognition of CA19-9, we hypothesized that optimizing the interactions observed in its holo-state structure would improve CA19-9 binding affinity.

AbLIFT starts by manual selection of amino acid positions at the light-heavy chain interface, followed by a completely automated three-step workflow: First, mutations that are rarely observed in a multiple-sequence alignment of the antibody’s homologs are eliminated at each position; second, each retained point mutation is atomically modeled in Rosetta and relaxed using an all-atom energy function (talaris14) which is dominated by van der Waals contacts, hydrogen bonding and implicit solvation [28], and highly destabilizing mutations are eliminated; last, all combinations of point mutations selected in the previous steps are enumerated in Rosetta, relaxed, and ranked by the all-atom energy function. These multipoint mutants are clustered, removing sequences that are fewer than three mutations from one another, and the top-ranked designs are screened experimentally.

We applied AbLIFT to the coordinates of 1116NS19.9 bound to CA19-9 and selected 17 variants that were calculated to have the most significant favorable change in Rosetta free energy (ΔΔ*G*) of the variable domain. Based on visual inspection, we chose heavychain positions Asp35, Thr93, Thr94, and Tyr98 (Fig. 7a) and light chain residues Ser43, Asp56, Tyr87, and Phe98 (Fig. 7b) for design. In a preliminary screen, we experimentally expressed all 17 designs and tested their binding using surface plasmon resonance (SPR) to a monomeric CA19-9 as an analyte in steady-state binding experiments with five-point concentration series ranging from 0.488 μM to 125 μM. These assays reflect the true microscopic affinities in the absence of avidity effects. Out of 17 variants, 15 either did not bind CA19-9 at all, or bound but displayed affinities that were weaker or similar to 1116NS19.9. Two designs displayed better binding affinities to CA19-9, of which we selected one called AbLIFT-15 for further analysis. Using steady-state analyses and an extended 14-points concentration series ranging from 500 μM to 0 μM of CA19-9, we measured for AbLIFT-15 a *K_D_* value of 1.7 μM, compared to a *K_D_* value of 14.7 μM for 1116NS19.9 (Fig. 7c), which represents a nearly tenfold improvement in affinity. We further measured the affinity of 5b1 to CA19-9 as a reference in this avidity-free system and determined the *K_D_* value to be 12.8 μM. Hence, AbLIFT-15 exhibits tenfold higher affinity compared to both 1116NS19.9 and 5b1 anti-CA19-9 antibodies.

**Fig 7.**
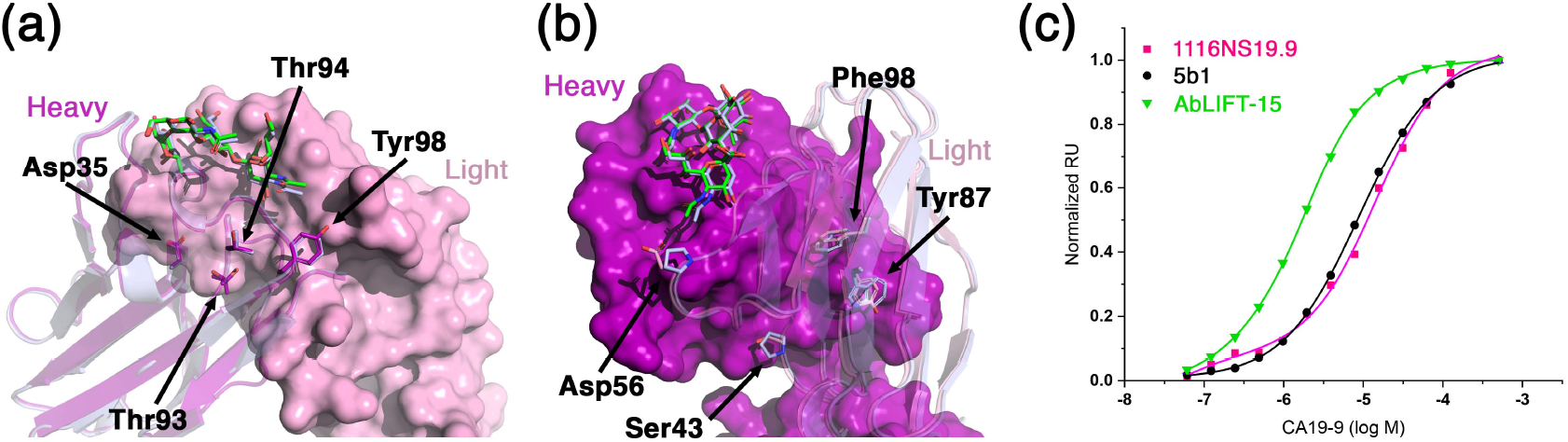
Recognition of CA19-9 and enhanced binding affinity of AbLIFT-15. (a) Superimposition of 1116NS19.9 (purple and pink for the heavy and light chains, respectively) and of AbLIFT-15 (pale cyan). The light chain is shown using surface representation, and the heavy chains are shown as ribbons. The four heavy-chain residues that were designed by the AbLIFT design protocol are indicated. (b) Superimposition of 1116NS19.9 (purple and pink for the heavy and light chains, respectively) and of AbLIFT-15 (pale cyan). The heavy chain is shown using surface representation, and the light chains are shown as ribbons. The four light-chain residues that were indicated by the AbLIFT design protocol are indicated. (c) Steady-state SPR analyses of 1116NS19.9, 5b1, and AbLIFT-15 using CA19-9 in a 2-fold dilution series starting at 500 μM. Binding experiments were repeated three times, and representative results are shown.

AbLIFT-15 has a total of seven mutations compared to 1116NS19.9, a large number of core mutations relative to mutants obtained in conventional antibody engineering and design efforts (T93A, T94V, and Y98F in the heavy chain and S43P, D56P, Y87W, and F98W in the light chain). To verify that the molecular recognition of CA19-9 by the designed antibody did not differ substantially from the parental antibody, we determined the bound complex structure at 1.4 Å resolution (Table 1). Remarkably, despite seven core mutations, the structure reveals an almost identical main-chain structure (RMSD of 0.22 Å for 220 Cα atoms of the variable regions) (Fig. 7a & 7b). Crucially, binding to CA19-9 was very similar to the conformation observed in the parental antibody 1116NS19.9 as predicted by the design model.

## DISCUSSION

In comparison to proteins, glycans are considered as more challenging target immunogens for the humoral immune system. The low immunogenicity of aberrant glycans is due in part to their potential flexibility and their highly hydrophilic nature. For this reason, understanding the molecular details of how antibodies bind to glycans may provide important biophysical insights for designing next-generation diagnostics and therapeutics. Here, we provide structural information for the recognition of CA19-9 by two different mAbs. Both 1116NS19.9 and 5b1 bind CA19-9 using extensive polar-interaction networks, which allow them to bind this small antigen with a *K_D_* of 14.7 μM and 12.8 μM, respectively. Despite being isolated from different hosts (*i.e.,* mouse and human, respectively) and against different targets, 1116NS19.9 [20,21] and 5b1 [22] recognize an almost identical conformation of CA19-9. This observation strongly implies that the configuration of CA19-9, as determined here, represents a preferred low-energy state of this carbohydrate antigen. These findings further suggest that the binding of antibodies to CA19-9 may be partially restricted in cases where the antigen cannot freely rotate with respect to the protein-surface that it modifies. In such scenarios, using a combination of antibodies like 1116NS19.9 and 5b1 may provide a more complete detection of CA19-9 than using each of the mAbs alone. This could be advantageous for either therapy or diagnostics.

Being relatively small and hydrophilic makes CA19-9 a challenging target for molecular binding and for conventional optimization strategies that target the complementarity-determining region for mutation. We previously used library screening to identify variants of 1116NS19.9 that can bind to CA19-9 more tightly [18]. In this current study, we extended our efforts and used structural data together with innovative computational design to produce AbLIFT-15. All the seven mutations that we have introduced to AbLIFT-15 are at the interface between the V_H_ and V_L_ and do not directly contact CA19-9. These results validate the use of AbLIFT to automatically, effectively and through a modest experimental effort enhance antibodies that have been mostly recalcitrant to conventional optimization approaches.

The shortcomings of CA19-9 as a marker for early diagnosis and screening for pancreatic cancer include false-positive results for patients with a benign pancreatic disease [10–12,29], and prompted the search of other biomarkers [30]. However, recent research demonstrates CA19-9-induced pancreatitis as a driving force for pancreatic cancer in a mouse model, justifying pancreatic-cancer monitoring for individuals with pancreatitis [14]. False-negative results are reported for 5-10% of the population [29]. In addition to the false positive and negative sub-populations, the sensitivity and specificity of the CA19-9 screening are not sufficient [10–12,31]. While false positive and negative results due to benign illness and genotpscreening are not sufficientye cannot be eliminated through the application of enhanced antibodies towards CA19-9, tighter binding may improve the positive predictive score for screening tests. Future research will be directed to studying whether the enhanced anti-CA19-9 antibodies provide a benefit in earlier and more accurate diagnosis for pancreatic cancer, either by detecting CA19-9 alone or in combination with other biomarkers [31]. In addition, achieving stronger binding to CA19-9 could facilitate the use of anti-CA19-9 antibodies as immunotherapeutic agents.

## MATERIALS AND METHODS

### Cloning of antibodies into yeast surface display (YSD) system and functional assay

Sequences of the variable domains (V_H_ and V_L_) of the anti-SLe^a^ mouse antibody 1116NS19.9 [20] and the human antibody 5b1 [22] were used to design scFv of (N’-V_H_)–(GGGGSGGGGSGGGGS linker)-(C’-V_L_), and DNA fragments synthesized by Integrated DNA Technologies Inc. (IDT, Israel). The scFv DNA sequence was optimized for codon usage compatible with expression in human cells, without altering the amino acid sequence. In addition, the scFv sequence was flanked by plasmid homology regions at the 5′ and 3′ ends (36 and 45 nucleotides, respectively). The flanking regions contained *5′-NdeI* and *3′-BamHI* restriction enzyme cloning site inframe with the scFv. Then, EBY100 yeast cells were transformed with each synthesized scFv and *NdeI BamHI* digested plasmid for *in vivo* ligation, as described [18]. The resulting scFv contained N’ HA and C’ c-Myc tags (encoded in the plasmid) that allowed to monitor surface expression.

### Induction of scFv expression on YSD system

To obtain scFv surface expression on yeast cells, 1116NS19.9-scFv-pETCON2 or 5b1-scFv-pETCON2 transfected yeast cells were cultured in SD-Trp a synthetic defined media (SD) lacking Tryptophan (Trp) [2% glucose (Sigma), 0.67% yeast nitrogen base w/o amino acids (BD), 0.54% Na_2_HPO_4_ (Sigma), 0.86% NaH_2_PO_4_ (Sigma) and 0.192% yeast synthetic drop-out medium supplements without Trp (Sigma)] at 30 °C, passaged 1:10 each day for three days, then scFv was expressed by changing the media to SG-Trp a synthetic galactose (SG) based media [2% galactose (Sigma), 0.2% glucose, 0.67% yeast nitrogen base w/o amino acids, 0.54% Na_2_HPO_4_, 0.86% NaH_2_PO_4_, and 0.192% yeast synthetic drop-out medium supplements without Trp] and the temperature to 20 °C, and cells were grown overnight to obtain scFv-YSD cells.

### Assessment of scFv functional reactivity by Fluorescence-Activated Cell Sorting (FACS)

Induced scFv-YSD cells were functionally analyzed for antigen binding by FACS, as described [18], with some modifications. Briefly, we used target antigens in a nanoparticle expression mode with multivalent expression on ~30 kDa polyacrylamide polymers carrying biotin tags (PAA-Bio; biotin ~every 5th amide group with 7–9 glycans per particle). Thus, we used polyvalent SLe^a^-PAA-Bio glycans nanoparticles. The non-specific target antigen Le^a^-PAA-Bio was used as a negative control. 5 × 10^6^ 1116NS19.9-scFv-expressing yeast cells or 5b1-scFv-expressing yeast cells were washed with 1 ml assay buffer (PBS, 0.5% ovalbumin) then incubated with 1 μM SLe^a^-PAA-biotin antigen and 1:50 diluted mouse-anti-c-Myc (4 μg/ml), both in 50 μl assay buffer, and for negative control, cells were incubated with 50 μl assay buffer, then all incubated for 1 h at room temperature (RT) with rotation. Cells were washed with 1 ml ice cold assay buffer, then incubated for 40 min on ice with APC-streptavidin and Alexa-Fluor-488-goat-anti-mouse IgG1 diluted 1:50 (10 μg/ml) and 1:200 (10 μg/ml) respectively in 50 μl assay buffer. Cells were washed with 1 ml ice cold PBS, then resuspended in 500 μl PBS. Cell fluorescence was measured by FACSort flow cytometry (Becton Dickinson) and analyzed with Kaluza analysis software. Double positive (APC-Ag^+^AF488-Ab^+^) yeast cells exemplified functional cloned scFv constructs (1116NS19.9-scFv and 5b1-scFv).

### Cloning and expression of antibodies as IgGs

Cloning was done by Gibson assembly as described [18], with some modifications. Variable heavy and light fragments of 1116NS19.9 or 5b1 were amplified by PCR. Reaction was made in Q5 reaction buffer, with 1 μl of plasmid DNA template (65–98 ng), 200 μM each dNTP, 1 U Q5 hot start high fidelity DNA polymerase (New England Biolabs), 500 nM each primer (Table 2 primers #1-4 for 1116NS19.9 or primers #5-8 for 5b1) complete volume to 50 μl with PCR grade water. PCR conditions were 95 °C for 2 min followed by 30 cycles of 95 °C for 30 s, 61 °C for 60 s, 72 °C for 60 s, and final incubation of 72 °C for 5 min. To remove template segments, the PCR product was supplemented with 6 μl of 10× CutSmart Buffer, 20 U DpnI (New England Biolabs), and completed the volume to 60 μl with PCR grade water, then incubated at 37 °C for 1 h. PCR digested fragments were purified from agarose gel by Zymoclean Gel DNA Recovery Kit (Zymo Research). Heavy and light chain full IgG p3BNC expression plasmids were divided to three parts for PCR amplification, variable region, left and right arms. Left and right arms of heavy and light p3BNC plasmids were amplified, digested and purified as described for variable regions using relevant primers (Table 2; primers #9-14). Of each fragment, variable region, right and left arms, 25 ng were taken for Gibson assembly. Reaction was made in isothermal reaction buffer containing 5% PEG 8000, 100 mM Tris-HCl pH 7.5, 10 mM MgCl2, 10 mM DTT, 0.2 mM of each dNTP and 10 mM NAD. To this buffer we added 0.04 U T5 exonuclease (NEB), 0.25U Phusion polymerase (NEB) and 40 U Taq DNA ligase (NEB), and ligation was made at 50 °C for 1 h. Plasmids were electroporated into XL1 Escherichia. coli, to validate the sequence and producing high amount of p3BNC expression plasmids. Human embryonic kidney 293A cells were then used to produce full length whole Ab 1116NS19.9 or Ab 5b1 from their respective p3BNC expression plasmids that were transfected with polyethylenimine reagent (PEI; Polysciences). Functionality of cloned secreted 1116NS19.9-hIgG1 and 5b1-hIgG1 antibodies was tested by ELISA against antigen-coated plates.

**Table 2.**
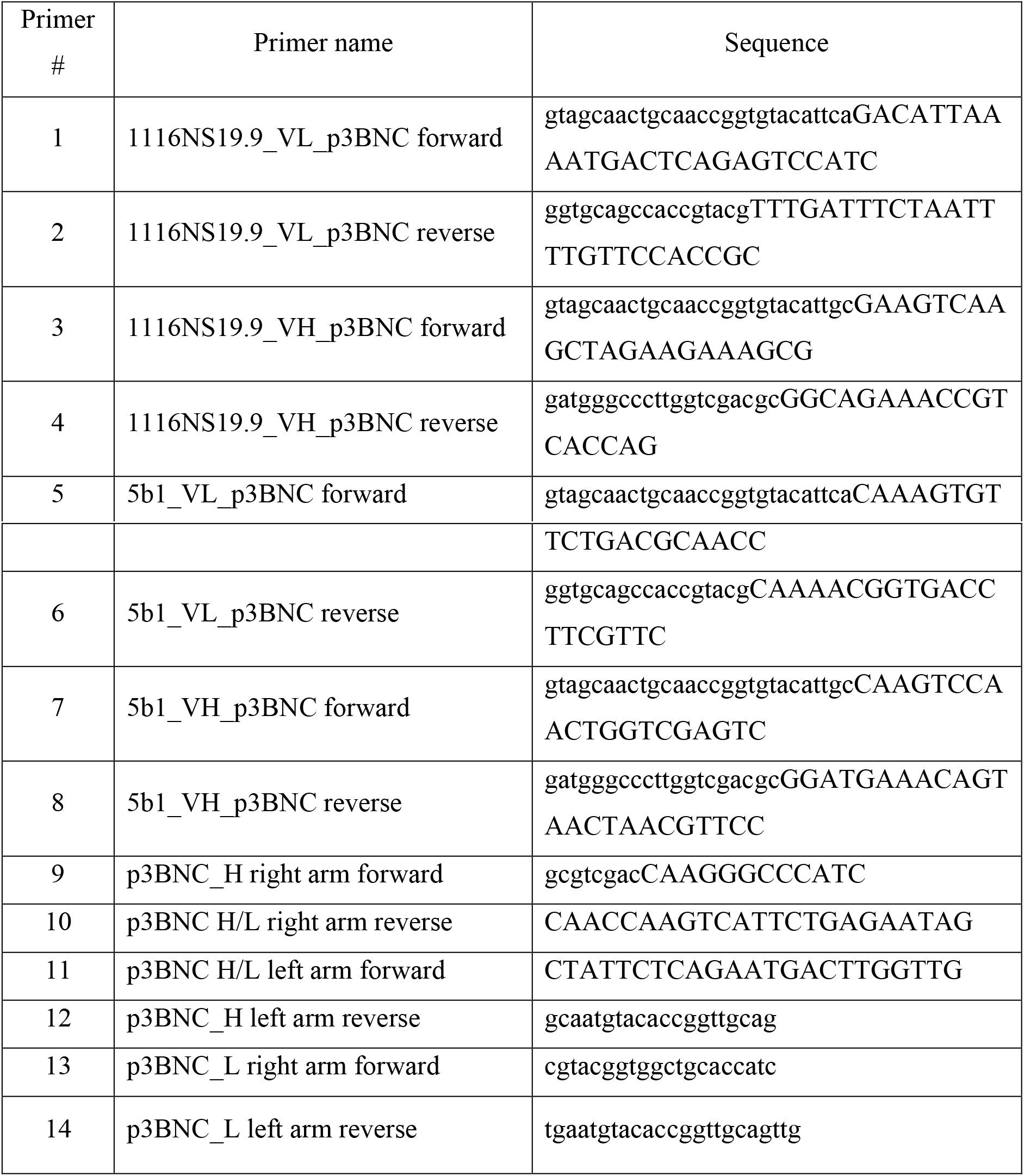

### Antibodies specificity by ELISA

Specificity was examined by binding of full-length 1116NS19.9-hIgG1 and 5b1-hIgG1 antibodies to various glycans by ELISA assay. 96-wells plate was coated with SLe^a^-PAA-Bio, Le^a^-PAA-Bio or PAA-Bio (GlycoTech) in duplicates at 0.25 μg/well overnight at 4 °C. Wells were blocked with blocking buffer (PBS pH7.4, 1% ovalbumin) for 1 hour at RT. Blocking buffer was removed and supernatant of 293A transfected cells containing 1116NS19.9-hIgG1 or 5b1-hIgG1 antibodies diluted 1:100 in blocking buffer was added at 100 μl/well, then incubated for two hours at RT. Plates were washed three times with PBST (PBS pH 7.4, 0.1% Tween), then incubated for 1 h at RT with HRP-goat-anti-human IgG 0.11 μg/ml in 100 μl PBS. After washing three times with PBST, wells were developed with 140 μl of O-phenylenediamine in 100 mM citrate-PO4 buffer, pH 5.5, and the reaction stopped with 40 μl of H_2_SO_4_ (4 M). Absorbance was measured at 490 nm on SpectraMax M3 (Molecular Devices). Specific binding was defined by subtracting the background readings obtained with the secondary antibody only.

### Synthesis of SLe^a^βProNH_2_

The tumor-associated carbohydrate antigen SLe^a^ in the form of SLe^a^βProN_3_ [Neu5Acα2–3Galβ1–3(Fucα1–4)GlcNAcβO(CH_2_)_2_CH_2_N_3_] was synthesized as previously described [25]. It was used to synthesize SLe^a^βProNH_2_ by catalytic hydrogenation as described below. To a stirred solution of SLe^a^βProN_3_ (5 mg) in water-methanol solution (2.1 ml, 1:2 by volume) in a round bottom flask (50 ml), 10% pallidum on charcoal Pd/C (2 mg) was added. The mixture was stirred under a hydrogen environment for 2 h. The solution was then passed through a filter to remove the catalyst. The solvent was removed under vacuum and the residue was dissolved in 0.5 ml of deionized water, frozen, and lyophilized to produce SLe^a^ProNH_2_ as a white powder.

### Protein expression and purification

To produce large amount of recombinant Abs for crystallization, we transfected HEK293F cells maintained in FreeSyle medium (Gibco) with the p3BNC plasmids encoding the heavy and the light chains of Abs. As a transfection reagent, we used 40 kDa polyethyleneimine (PEI) (Polysciences) with a DNA / polyethyleneimine ratio of 1 μg / 3 μl with a total of 1 mg DNA per 1 L of cells at 1M cells/ml. Cells were maintained for 5-7 days in suspension before harvesting the supernatants. After clarifying the supernatants by centrifugation, Abs were captured using protein-A affinity chromatography (GE Lifesciences). Abs were eluted using 0.1 M citric acid pH 3.0 buffer, which was subsequently adjusted to pH 8.0 using Tris-HCl. For obtaining the Fab portions, papain enzyme (Sigma-Aldrich) was used to digest Abs in enzyme and protein ratio being ~1:80. Cutting buffer contained 20 mM Cysteine-HCl (Sigma-Aldrich) and 10 mM EDTA tittered to pH 7.0 with Tris buffer pH 8.0. Cutting was performed for 90 minutes in 37° C. Negative protein-A was performed to remove Fc fragments followed by SEC on a Superdex200 10/300 column (GE Lifesciences).

### Crystallization

For protein crystallization, we used a mosquito crystallization robot (TTP Labtech) to set vapor diffusion in sitting drop experiments using 96-well iQ plates (TTP Labtech). At each well, we tested three ratios of protein (80, 120, and 160 nl) to reservoir (120 nl). PEGrx-HT screen (Hampton Research) was used to identify initial hits for apo-Ab 1116NS19.9 for the condition containing 0.10% w/v n-Octyl-b-D-glucoside, 0.1 M Sodium citrate tribasic dihydrate pH 5.5, and 22% w/v Polyethylene glycol 3,350. Further optimization was done by growing the crystals in 7.5% ethylene glycol for cryo-preservation. Protein with ligand CA19-9 was mixed in a ratio of 1:1.2 protein to CA19-9, protein samples gave crystals when grown in 24-well sitting plates with a 1:1 ratio of protein to reservoir. Ab 5b1 apo- and halo-protein crystals were crystallized using the same reservoir conditions containing 0.1M NaCl, 0.1M bis-tris propane pH 9.0, 18% polyethylene glycol 1,500 and 5% glycerol. The protein to CA19-9 ratio was 1:1, and the protein to reservoir ratio was 1.75:1. All crystals were grown at 20° C.

### Data collection, structure solution and refinement

X-ray diffraction data were collected at the European Synchrotron Radiation Facility (ESRF) using a Pilatus 6M detector at 100° K. Data up to 1.5 Å at beamline ID23-1 was collected for the apo and halo Fab 1116NS19.9 belonging to the tetragonal and orthorhombic space groups, respectively. Data were indexed, integrated, and scaled using XDS [32]. We used Phaser [33] to obtain a molecular replacement solution with the structure of NIH45-46 (PDB: 3u7w) and used the solved structure for molecular replacement of Fab 5b1 halo- and apo-proteins. Data for halo- and apo-Ab 5b1 were collected at beamline ID23-2 using a Pilatus 2M detector at resolutions of 2.4 Å for the apo and halo structures, which belong to the hexagonal space group with 3 and 6 molecules in the asymmetric unit, respectively. All models were manually traced into electron density maps using Coot [34] and refined using Phenix Refine [35] in an iterative fashion.

### Surface Plasmon Resonance (SPR) measurements

All measurements were performed using a Biacore T200 (GE Healthcare) at 25 °C. Abs were immobilized to a protein-A chip from a stock of 20 μg/ml at similar surface densities. CA19-9 glycan in TBS buffer with azide 0.02% was used as an analyte in a series of increasing concentrations ranging from 0.488 μM to 125 μM using five steps for preliminary screening and ranging from 0 until 500 μM with 14 steps total for in-depth analyses. Affinity constants were calculated by measuring binding levels at steady states and fitting binding curves to the data using the Biacore T200 evaluation software. We used washing with binding buffer without a regeneration step for achieving baseline signal after each injection. The protein-A sensor chip was eventually regenerated using 10 mM glycine–HCl, pH 1.5.

### Structure analysis and representation

Analyses and structural figures were generated using PyMol. Buried surface area calculations were performed using the AreaMol tool in the CCP4 program suite [36].

### AbLIFT design

AbLIFT was applied essentially as described [19]. Briefly, starting from the structure of 1116NS19.9, we manually selected eight positions at the interface between the variable light and heavy chains for design calculations. A multiple sequence alignment of all homologs was obtained using default parameters and a position-specific scoring matrix was computed using PSI-BLAST [37]. At each position, mutations that exhibited a PSSM score < −1 were eliminated from the design sequence choices. Next, each remaining mutation was modeled using the Rosetta atomistic modelling and design package [38] and relaxed using the talaris14 energy function [28], which is dominated by van der Waals contacts, hydrogen bonding and solvation. Mutations that exhibited a total energy > +1 Rosetta energy units higher than the relaxed structure of the parental antibody were further eliminated. With the remaining identities, 130,111 combinations of mutations at 8 positions that were at least 3 mutations different from the parental antibody were modeled and relaxed in Rosetta. The mutants were then ranked according to their all-atom energy, clustered by requiring that each multipoint mutant exhibit at least 3 mutations relative to any other, and the top 17 designs were selected for experimental analysis.

### Accession numbers

Coordinates and structure functions were deposited in the protein data bank under the accession codes that are listed in Table 1.

## Author contribution

**Aliza Borenstein-Katz**: Investigation, Formal analysis, Writing Original-draft, Visualization. **Shira Warszawski**: Software, Formal analysis, Writing Original-draft. **Ron Amon**: Investigation, Resources, Writing Original-draft, Visualization. **Nova Tasnima**: Resources. **Hai Yu**: Resources. **Xi Chen**: Resources. **Vered Padler-Karavani**: Conceptualization, Investigation, Writing Review & Editing, Visualization, Formal analysis, Supervision, Funding acquisition. **Sarel Jacob Fleishman**: Conceptualization, Investigation, Writing Review & Editing, Software, Formal analysis, Supervision, Resources. **Ron Diskin**: Conceptualization, Investigation, Writing Review & Editing, Formal analysis, Visualization, Supervision.

## ACKNOWLEDGEMENTS

We are grateful to Royant Antoine and Zubieta Chloe at the ESRF, Grenoble, France, for assisting in using beamlines ID23-1 and ID23-2. The Diskin lab is supported by research grants from the Ernst I Ascher foundation, Ben B. and Joyce E. Eisenberg Foundation, Estate of Emile Mimran, Jeanne and Joseph Nissim Center for Life Sciences Research, Dov and Ziva Rabinovich Endowed Fund for Structural Biology, Donald Rivin, Stanley and Tanya Rossby Endowment Fund, Natan Sharansky, Dr. Barry Sherman Institute for Medicinal Chemistry, as well as from the Israel Science Foundation (grants No. 3147/19, 209/20, & 682/16). Research in the Fleishman lab was supported by a Consolidator Award from the European Research Council (815379) and by a charitable donation in memory of Sam Switzer. Research in the Padler-Karavani lab was funded by the European Union H2020 Program grants (ERC-2016-STG-716220), and UC Davis – Israel Collaborations in Research Proposal award UC Davis (to V.P-K and X.C.).

## REFERENCES

[1] J.R. Kintzing, M. V. Filsinger Interrante, J.R. Cochran, Emerging Strategies for Developing Next-Generation Protein Therapeutics for Cancer Treatment, Trends Pharmacol. Sci. (2016). https://doi.org/10.1016/j.tips.2016.10.005.

[2] S.R. Stowell, T. Ju, R.D. Cummings, Protein glycosylation in cancer, Annu. Rev. Pathol. Mech. Dis. (2015). https://doi.org/10.1146/annurev-pathol-012414-040438.

[3] A. Varki, R. Kannagi, B. Toole, P. Stanley, Glycosylation Changes in Cancer, 2015.

[4] V. Padler-Karavani, Aiming at the sweet side of cancer: Aberrant glycosylation as possible target for personalized-medicine, Cancer Lett. 352 (2014) 102–112. https://doi.org/10.1016/j.canlet.2013.10.005.

[5] S.S. Pinho, C.A. Reis, Glycosylation in cancer: mechanisms and clinical implications, Nat. Rev. Cancer. 15 (2015) 540–555. https://doi.org/10.1038/nrc3982.

[6] K.F. Boligan, C. Mesa, L.E. Fernandez, S. Von Gunten, Cancer intelligence acquired (CIA): Tumor glycosylation and sialylation codes dismantling antitumor defense, Cell. Mol. Life Sci. (2015). https://doi.org/10.1007/s00018-014-1799-5.

[7] R. Kannagi, Carbohydrate antigen sialyl Lewis a - Its pathophysiological significance and induction mechanism in cancer progression, Chang Gung Med. J. (2007).

[8] R. Amon, E.M. Reuven, S. Leviatan Ben-Arye, V. Padler-Karavani, Glycans in immune recognition and response, Carbohydr. Res. (2014). https://doi.org/10.1016/j.carres.2014.02.004.

[9] M. Ugorski, A. Laskowska, Sialyl Lewis a: A tumor-associated carbohydrate antigen involved in adhesion and metastatic potential of cancer cells, Acta Biochim. Pol. 49 (2002) 303–311.

[10] U.K. Ballehaninna, R.S. Chamberlain, The clinical utility of serum CA 19-9 in the diagnosis, prognosis and management of pancreatic adenocarcinoma: An evidence based appraisal, J. Gastrointest. Oncol. 3 (2012) 105–119. https://doi.org/10.3978/j.issn.2078-6891.2011.021.

[11] Z. Huang, F. Liu, Diagnostic value of serum carbohydrate antigen 19-9 in pancreatic cancer: a meta-analysis, Tumor Biol. (2014). https://doi.org/10.1007/s13277-014-1995-9.

[12] R. Passerini, M.C. Cassatella, S. Boveri, M. Salvatici, D. Radice, L. Zorzino, C. Galli, M.T. Sandri, The pitfalls of CA19-9: Routine testing and comparison of two automated immunoassays in a reference oncology center, Am. J. Clin. Pathol. (2012). https://doi.org/10.1309/AJCPOPNPLLCYR07H.

[13] S. Bussom, M.W. Saif, Methods and rationale for the early detection of pancreatic cancer, in: J. Pancreas, 2010.

[14] D.D. Engle, H. Tiriac, K.D. Rivera, A. Pommier, S. Whalen, T.E. Oni, B. Alagesan, E.J. Lee, M.A. Yao, M.S. Lucito, B. Spielman, B. Da Silva, C. Schoepfer, K. Wright, B. Creighton, L. Afinowicz, K.H. Yu, R. Grützmann, D. Aust, P.A. Gimotty, K.S. Pollard, R.H. Hruban, M.G. Goggins, C. Pilarsky, Y. Park, D.J. Pappin, M.A. Hollingsworth, D.A. Tuveson, The glycan CA19-9 promotes pancreatitis and pancreatic cancer in mice, Science (80-.). 364 (2019) 1156–1162. https://doi.org/10.1126/science.aaw3145.

[15] J.C. Manimala, T.A. Roach, Z. Li, J.C. Gildersleeve, High-throughput carbohydrate microarray profiling of 27 antibodies demonstrates widespread specificity problems, Glycobiology. (2007). https://doi.org/10.1093/glycob/cwm047.

[16] E. Sterner, N. Flanagan, J.C. Gildersleeve, Perspectives on Anti-Glycan Antibodies Gleaned from Development of a Community Resource Database, ACS Chem. Biol. (2016). https://doi.org/10.1021/acschembio.6b00244.

[17] R. Amon, O.C. Grant, S. Leviatan Ben-Arye, S. Makeneni, A.K. Nivedha, T. Marshanski, C. Norn, H. Yu, J.N. Glushka, S.J. Fleishman, X. Chen, R.J. Woods, V. Padler-Karavani, A combined computational-experimental approach to define the structural origin of antibody recognition of sialyl-Tn, a tumor-associated carbohydrate antigen, Sci. Rep. (2018). https://doi.org/10.1038/s41598-018-29209-9.

[18] R. Amon, R. Rosenfeld, S. Perlmutter, O.C. Grant, S. Yehuda, A. Borenstein-Katz, R. Alcalay, T. Marshanski, H. Yu, R. Diskin, R.J. Woods, X. Chen, V. Padler-Karavani, Directed evolution of therapeutic antibodies targeting glycosylation in cancer, Cancers (Basel). (2020). https://doi.org/10.3390/cancers12102824.

[19] M.S. and S.J.F. Shira Warszawski, Aliza Katz, Lev Khmelnitsky, Gili Ben Nissan, Rosalie Lipsh, Gabriel Javitt, Orly Dym, Tamar Unger, Orli Knop, Shira Albeck, Ron Diskin, Deborah Fass, Optimizing antibody affinity and stability by the automated design of the variable light-heavy chain interfaces, PLOS Comput. Biol. (2019) 1–24.

[20] H. Koprowski, Z. Steplewski, K. Mitchell, M. Herlyn, D. Herlyn, P. Fuhrer, Colorectal carcinoma antigens detected by hybridoma antibodies, Somatic Cell Genet. 5 (1979) 957–971. https://doi.org/10.1007/BF01542654.

[21] R. Passerini, D. Riggio, M. Salvatici, L. Zorzino, D. Radice, M.T. Sandri, Interchangeability of measurements of CA 19-9 in serum with four frequently used assays: An update, Clin. Chem. Lab. Med. 45 (2007) 100–104. https://doi.org/10.1515/CCLM.2007.003.

[22] R. Sawada, S.M. Sun, X. Wu, F. Hong, G. Ragupathi, P.O. Livingston, W.W. Scholz, Human monoclonal antibodies to sialyl-Lewis a (CA19.9) with potent CDC, ADCC, and antitumor activity, Clin. Cancer Res. 17 (2011) 1024–1032. https://doi.org/10.1158/1078-0432.CCR-10-2640.

[23] J.L. Houghton, D. Abdel-Atti, W.W. Scholz, J.S. Lewis, Preloading with Unlabeled CA19.9 Targeted Human Monoclonal Antibody Leads to Improved PET Imaging with 89Zr-5B1, Mol. Pharm. 14 (2017) 908–915. https://doi.org/10.1021/acs.molpharmaceut.6b01130.

[24] G. Chao, W.L. Lau, B.J. Hackel, S.L. Sazinsky, S.M. Lippow, K.D. Wittrup, Isolating and engineering human antibodies using yeast surface display, Nat. Protoc. (2006). https://doi.org/10.1038/nprot.2006.94.

[25] N. Tasnima, H. Yu, X. Yan, W. Li, A. Xiao, X. Chen, Facile chemoenzymatic synthesis of Lewis a (Lea) antigen in gram-scale and sialyl Lewis a (sLea) antigens containing diverse sialic acid forms, Carbohydr. Res. (2019). https://doi.org/10.1016/j.carres.2018.12.004.

[26] S. Re, W. Nishima, N. Miyashita, Y. Sugita, Conformational flexibility of N-glycans in solution studied by REMD simulations, Biophys. Rev. 4 (2012) 179–187. https://doi.org/10.1007/s12551-012-0090-y.

[27] A.C. Wallace, R.A. Laskowski, J.M. Thornton, LIGPLOT: a program to generate schematic diagrams of protein-ligand interactions The LIGPLOT program automatically generates schematic 2-D representations of proteinligand complexes from standard Protein Data Bank file input, Protein Eng. (1995).

[28] M.J. O’Meara, A. Leaver-Fay, M.D. Tyka, A. Stein, K. Houlihan, F. Dimaio, P. Bradley, T. Kortemme, D. Baker, J. Snoeyink, B. Kuhlman, Combined covalent-electrostatic model of hydrogen bonding improves structure prediction with Rosetta, J. Chem. Theory Comput. (2015). https://doi.org/10.1021/ct500864r.

[29] M.J. Duffy, C. Sturgeon, R. Lamerz, C. Haglund, V.L. Holubec, R. Klapdor, A. Nicolini, O. Topolcan, V. Heinemann, Tumor markers in pancreatic cancer: A European Group on Tumor Markers (EGTM) status report, Ann. Oncol. 21 (2009) 441–447. https://doi.org/10.1093/annonc/mdp332.

[30] B. Staal, Y. Liu, D. Barnett, P. Hsueh, Z. He, C.F. Gao, K. Partyka, M.W. Hurd, A.D. Singhi, R.R. Drake, Y. Huang, A. Maitra, R.E. Brand, B.B. Haab, The Stra plasma biomarker: Blinded validation of improved accuracy over CA19-9 in pancreatic cancer diagnosis, Clin. Cancer Res. (2019). https://doi.org/10.1158/1078-0432.CCR-18-3310.

[31] D. Dong, L. Jia, L. Zhang, N. Ma, A. Zhang, Y. Zhou, L. Ren, Periostin and CA242 as potential diagnostic serum biomarkers complementing CA19.9 in detecting pancreatic cancer, Cancer Sci. 109 (2018) 2841–2851. https://doi.org/10.1111/cas.13712.

[32] W. Kabsch, Integration, scaling, space-group assignment and post-refinement, Acta Crystallogr. Sect. D. 66 (2010) 133–144. https://doi.org/10.1107/S0907444909047374.

[33] A.J. McCoy, R.W. Grosse-Kunstleve, P.D. Adams, M.D. Winn, L.C. Storoni, R.J. Read, Phaser crystallographic software, J. Appl. Crystallogr. 40 (2007) 658–674. https://doi.org/10.1107/S0021889807021206.

[34] P. Emsley, B. Lohkamp, W.G. Scott, K. Cowtan, Features and development of Coot, Acta Crystallogr. Sect. D Biol. Crystallogr. 66 (2010) 486–501. https://doi.org/10.1107/S0907444910007493.

[35] P.D. Adams, P. V. Afonine, G. Bunkóczi, V.B. Chen, I.W. Davis, N. Echols, J.J. Headd, L.W. Hung, G.J. Kapral, R.W. Grosse-Kunstleve, A.J. McCoy, N.W. Moriarty, R. Oeffner, R.J. Read, D.C. Richardson, J.S. Richardson, T.C. Terwilliger, P.H. Zwart, PHENIX: A comprehensive Python-based system for macromolecular structure solution, Acta Crystallogr. Sect. D Biol. Crystallogr. 66 (2010) 213–221. https://doi.org/10.1107/S0907444909052925.

[36] C.C. Project, The CCP4 suite: Programs for protein crystallography, Acta Crystallogr. Sect. D Biol. Crystallogr. 50 (1994) 760–763. https://doi.org/10.1107/S0907444994003112.

[37] S.F. Altschul, T.L. Madden, A.A. Schäffer, J. Zhang, Z. Zhang, W. Miller, D.J. Lipman, Gapped BLAST and PSI-BLAST: A new generation of protein database search programs, Nucleic Acids Res. (1997). https://doi.org/10.1093/nar/25.17.3389.

[38] A. Leaver-Fay, M. Tyka, S.M. Lewis, O.F. Lange, J. Thompson, R. Jacak, K. Kaufman, P.D. Renfrew, C.A. Smith, W. Sheffler, I.W. Davis, S. Cooper, A. Treuille, D.J. Mandell, F. Richter, Y.E.A. Ban, S.J. Fleishman, J.E. Corn, D.E. Kim, S. Lyskov, M. Berrondo, S. Mentzer, Z. Popović, J.J. Havranek, J. Karanicolas, R. Das, J. Meiler, T. Kortemme, J.J. Gray, B. Kuhlman, D. Baker, P. Bradley, Rosetta3: An object-oriented software suite for the simulation and design of macromolecules, in: Methods Enzymol., 2011. https://doi.org/10.1016/B978-0-12-381270-4.00019-6.

